# Genome sequence of the oyster mushroom *Pleurotus ostreatus* strain PC9

**DOI:** 10.1101/2020.09.03.281683

**Authors:** Yi-Yun Lee, Guillermo Vidal-Diez de Ulzurrun, Erich M. Schwarz, Jason E. Stajich, Yen-Ping Hsueh

## Abstract

The oyster mushroom *Pleurotus ostreatus* is a basidiomycete commonly found in the rotten wood and it is one of the most cultivated edible mushrooms globally. *P. ostreatus* is also a carnivorous fungus, which can paralyze and kill nematodes within minutes. However, the molecular mechanisms of the predator-prey interactions between *P. ostreatus* and nematodes remain unclear. PC9 and PC15 are two model strains of *P. ostreatus* and the genomes of both strains have been sequenced and deposited at the Joint Genome Institute (JGI). These two monokaryotic strains exhibit dramatic differences in growth, but because PC9 grows more robustly in laboratory conditions, it has become the strain of choice for many studies. Despite the fact that PC9 is the common strain for investigation, its genome is fragmentary and incomplete relative to that of PC15. To overcome this problem, we used PacBio long reads and Illumina sequencing to assemble and polish a more integrated genome for PC9. Our PC9 genome assembly, distributed across 17 scaffolds, is highly contiguous and includes six telomere-to-telomere scaffolds, dramatically improving the genome quality. We believe that our PC9 genome resource will be useful to the fungal research community investigating various aspects of *P. ostreatus* biology.

## Introduction

*Pleurotus ostreatus* is a common edible basidiomycete that ranks second for global mushroom consumption (Cohen et al. 2002; Sánchez 2010). In the wild, this fungus prefers temperate habitats, such as subtropical forest (Hilber 1997), where it extracts nutrients from dead or dying trees (Karim et al. 2016). In order to degrade the wood, *P. ostreatus* releases several enzymes such as lignin, cellulose, and hemicellulose (Sánchez 2009). Nematicidal toxins produced by *P. ostreatus* under starvation conditions, such as, trans-2-decenedioic, can paralyze and kill their nematode-prey (Barron and Thorn 1987; Kwok et al. 1992). Another potential toxin, linoleic acid, reduces nematodes head size (Satou et al. 2008). When nematodes contact the mycelium of *P. ostreatus*, fungal toxins enter the prey through their sensory cilia, leading to hypercontraction and calcium influx of pharyngeal and body wall muscles, ultimately causing necrosis process of the neuromuscular system (Lee et al. 2020). However, the identity of the toxins and how exactly *P. ostreatus* induces rapid cell necrosis remain unclear. The *P. ostreatus* dikaryotic strain N001 that produces fruiting bodies is a common commercial strain (Larraya et al. 1999a; Peñas et al. 1998). In Larraya et al. (1999b), differentiated two monokaryotic strains (PC9 and PC15) from N001 in order to study reproduction and cloning in this mushroom. These two monokaryotic strains of *P. ostreatus* exhibit differences in growth pattern, with PC9 exhibiting faster growth relative to PC15. Although the genomes of these two monokaryotic strains have already been sequenced (Riley et al. 2014; Alfaro et al. 2016; Castanera et al. 2016), the quality of PC9 genome is poor for current standards. Whereas the latest version of the PC15 genome comprises only 12 scaffolds, the currently available PC9 genome (hereafter denoted PC9_JGI) is still distributed across 572 scaffolds, and most of which size are smaller than 1-kb (Alfaro et al. 2016; Castanera et al. 2016). Despite PC15 having a more complete genome than PC9, PC9 is often the preferred strain for research due to its robust growth in the laboratory. For instance, PC9 has been used to study ligninolytic activity (Nakazawa et al. 2017b; Nakazawa et al. 2017a; Nakazawa et al. 2019), gene modification (Yoav et al. 2018), and transformation (Nakazawa et al. 2016). Genetic manipulation has proven to significantly increase the production of primary and secondary metabolites in several filamentous fungi that are important in industry (Banerjee et al. 2003; Kück and Hoff 2010). Therefore, a high-quality PC9 genome is needed to enable scientists to conduct advanced genomic studies and to facilitate functional analyses of *P. ostreatus* genes.

Consequently, we decided to sequence and annotate an updated genome for PC9. In this study, we used long PacBio reads and Illumina reads to assemble the genome of *Pleurotus ostreatus* strain PC9 and functionally annotated its predicted proteins. The final assembly, obtained by merging draft assemblies built with CANU (Koren et al.) and FALCON (Chin et al.), was distributed across 17 scaffolds, including six telomere-to-telomere scaffolds. In addition, the N50 of our assembly is ∼3.5 Mb, which ranks it among the best for currently available *Pleurotus* genomes. In summary, here we present a new high-quality PC9 genome that will be help the fungal research community to study this important mushroom.

## Materials and Methods

### Fungal strain and DNA extraction

*Pleurotus ostreatus* strain PC9 was cultured on yeast extract, malt extract and glucose medium (YMG) at 25 **°**C for 7 days. Mycelia were collected from original plates (5-cm) and added to a 100 mL YMG liquid medium shaken at 200 r.p.m. at 25 **°**C for 3 days. After 3 days, the culture was transferred to yeast nitrogen base without amino acids liquid medium (YNB), and it was shaken at 200 r.p.m. for 2 additional days. DNA was extracted by using cetyltrimethylammonium bromide (CTAB) and purified with chloroform, isopropanol, and phenol-chloroform.

### Genome sequencing and assembly

PacBio long reads were sequenced from the *Pleurotus ostreatus* PC9 genome using the Pacbio RSII platform, and a PacBio SMRTBell Template Prep Kit 1.0 SPv3 was used for library preparation. This approach resulted in a total of ∼0.3 M reads with a mean length of 14,742 base pairs (bp), representing 161X genome coverage for PC9. The Pacbio reads were used to construct two draft assemblies. The first assembly was built with Canu (v1.7) (Koren et al. 2017) using the parameters genomeSize=36m, useGrid=false, and maxThreads=8. The second assembly was built with Falcon (pbalign v0.02) (Chin et al. 2016). For this latter assembly, the configuration file was downloaded from: *https://pb-falcon.readthedocs.io/en/latest/parameters.html*, and the parameter Genome_size was set to 36 Mb since published *P. ostreatus* published genome sizes range from 34.3 to 35.6 Mb (Riley et al. 2014; Alfaro et al. 2016; Castanera et al. 2016). Next, we used Quiver (genomicconsensus v2.3.2) (Chin et al. 2013) to polish both these PacBio assemblies. Information on the raw polished draft assemblies is presented in Table S1.

The Canu assembly contained more telomeric regions, whereas the Falcon assembly was more contiguous. We merged both assemblies using Quickmerge (v0.3) (Chakraborty et al. 2016), with Canu as reference and Falcon as donor (Canu-Falcon) or *vice versa* (Falcon-Canu). The Falcon-Canu assembly showed better contiguity, having only 29 scaffolds (Table S2), so we selected it as the basis for our final assembly. Redundant contigs were detected using nucmer (Mummer4) (Marçais et al. 2018), which were subsequently filtered out of the assembly. Nucmer was also used to detect large overlapping regions within the scaffolds. When an overlapping region exceeded 10-kb, presented high identity (>99%), and lay at the end or beginning of the scaffolds, we manually merged the two scaffolds at the overlapping region (Table S3). In total, 10 scaffolds were merged in this way, yielding five larger and more complete scaffolds (scaffolds 2, 4, 6, 7, and 13 of the final assembly) (Table S3). Finally, the cleaned assembly consisting of 17 scaffolds was further polished using Illumina reads and Pilon (Walker et al. 2014). A total of 100 M Illumina sequence pair-end reads of 151 bp were used for Pilon polishing.

### Genome annotation

To annotate the assembled PC9 genome, we used *funannotate* (v1.5.2) (Jon Love et al. 2019) with the pipeline described in *https://funannotate.readthedocs.io/en/latest/tutorials.html*. We used the following commands: *funannotate* mask, to softmask the genome, *funannotate* training and *funannoate predict* to generate preliminary gene models and consensus gene models [using: AUGUSTUS (Stanke and Waack 2003), GeneMark (Borodovsky and McIninch 1993), and EVidenceModeler (Haas et al. 2008)], and *funannotate annotate* to add functional annotation.

### Genome analysis and comparison

General assembly statistics for example length and N50 of the scaffolds/contigs were calculated from the assembly fasta file using Perl scripts count_fasta_residues.pl (*https://github.com/SchwarzEM/ems_perl/blob/master/fasta/count_fasta_residues.pl*). BUSCO completeness was computed using BUSCO 3.0.1 (Simão et al. 2015; Waterhouse et al. 2018) against the Basidiomycota dataset basidiomycata_odb9 (Simão et al. 2015; Waterhouse et al. 2018). Repetitive elements were identified using a custom-made repeat library created according to a pipeline described previously (Coghlan et al. 2018). The repeat library was used as input for RepeatMasker (Smit 2013-2015), and the results were further analyzed using the one_code_to_find_them all script (Bailly-Bechet et al. 2014). The presence of telomeres in the scaffolds was established by searching for the telomeric repeats (TTAGGG)n (Pérez et al. 2009). Finally, we used Circos (v0.69.0) (Krzywinski et al. 2009) and D-genies (minimap v2) (Cabanettes and Klopp 2018) to illustrate the different genomic features of our assembly and to compare it to the previously published PC9_JGI genome (Alfaro et al. 2016; Castanera et al. 2016).

### Data availability

The final assembled and newly annotated genomes of *P. ostreatus* PC9 (denoted PC9_AS) has been uploaded to NCBI and is available with accession code: JACETU000000000.

## Results

### A new genome assembly of the *Pleurotus ostreatus* PC9 strain

Long PacBio and Illumina reads were used to sequence *P. ostreatus* strain PC9 to improve genome quality. The size of our *P. ostreatus* PC9_AS genome assembly is ∼35.0 Mb (Table 1), which concurs with the sizes of the currently available *P. ostreatus* genomes: PC9_JGI (35.6 Mb) and PC15 (34.3 Mb) (Riley et al. 2014; Alfaro et al. 2016; Castanera et al. 2016). A comparison with other *Pleurotus* species revealed that our *P. ostreatus* genome is smaller than that of *P. eryngii* strain 183 (43.8 Mb) (Yang et al. 2016), *P. tuoliensis* strain JKBL130LB (48.2 Mb) (Zhang et al. 2018), and *P. platypus* strain MG11 (40.0 Mb) (Li et al. 2018). PC9_AS is distributed across 17 scaffolds, with the maximum and minimum scaffold sizes being 4.86 Mb and 9.1 kb, respectively. The N50 value of our assembly data is 3.5 Mb, which ranks it highest among the available *Pleurotus* genomes, including for PC9_JGI (N50 = 2.09 Mb) (Alfaro et al. 2016; Castanera et al. 2016), PC15 (N50 = 3.27 Mb) (Riley et al. 2014; Alfaro et al. 2016; Castanera et al. 2016), CCMSSC03989 (N50 = 2.85 Mb) (Wang et al. 2018), *P. tuoliensis* strain JKBL130LB (N50 = 1.17 Mb) (Zhang et al. 2018), and *P. eryngii* strain 183 (N50 = 509 kb) (Yang et al. 2016). The total annotated gene number in PC9_AS is 11,875, which is slightly fewer than for PC9_JGI (12,206 genes). The completeness of our PC9 genome assembly was assessed with BUSCO (Simão et al. 2015; Waterhouse et al. 2018) using a Basidiomycota dataset. We obtained a 97.2% BUSCO completeness, with 1289 complete BUSCOs, 1284 complete and single copy BUSCOs (99.6%), 14 complete and duplicate BUSCOs (1.1%), 9 fragmented BUSCOs (0.7%), and 28 missing BUSCOs genes (2.2%). Overall, our statistical analysis suggests that our PC9_AS assembly is more completed and integrated than that of PC9_JGI (Alfaro et al. 2016; Castanera et al. 2016).

**Table 1.** Genomic features of the three *P. ostreatus* genome assemblies. nt, nucleotides.

| General features | PC9_AS | PC9_JGI | PC15 |
| --- | --- | --- | --- |
| Total nt | 35,032,978 | 35,630,309 | 34,342,730 |
| Number of scaffolds | 17 | 572 | 12 |
| N50 scaffold size, nt | 3,500,734 | 2,086,289 | 3,270,165 |
| N90 scaffold size, nt | 2,134,864 | 159,002 | 1,880,400 |
| Scaffold max. nt | 4,859,873 | 4,430,591 | 4,830,258 |
| Scaffold min. nt | 9,086 | 2,001 | 280,724 |
| Number of contigs | 17 | 3272 | 13 |
| N50 contig size, nt | 3,500,734 | 99,058 | 3,270,165 |
| N90 contig size, nt | 2,134,864 | 13,737 | 1,571,664 |
| GC content, % | 50.79 | 50.94 | 50.95 |
| Genes | 11,875 | 12,206 | 12,330 |
| BUSCO completeness, % | 97.2 | 97.2 | 97.6 |

The genome architecture of *Pleurotus ostreatus* strain PC9 is shown in Figure 1A. We used the highly conserved sequence (TTAGGG)n to determine telomere locations (Moyzis et al. 1988), which represent the ends of the chromosomes in fungi (Farman and Leong 1995; Schechtman 1990). Out of 17 scaffolds, six possess telomeric repeats at both ends, indicating that these scaffolds represent complete chromosomes (Figure 1A). In contrast, three scaffolds have a telomere at only one end, and the remaining scaffolds lack any apparent telomeric repeats. Interestingly, the small scaffolds 13 (93,126 bp) and 14 (62,249 bp) have telomeric repeats, so they may constitute the ends of other scaffolds lacking telomeres. Larraya et al. (1999b) used pulsed-field gel electrophoresis to determine how many chromosomes of PC9 and PC15 have and reported a total of eleven chromosomes in *P. ostreatus*, thus suggesting that our assembly is close to the chromosome level.

**Fig. 1.**
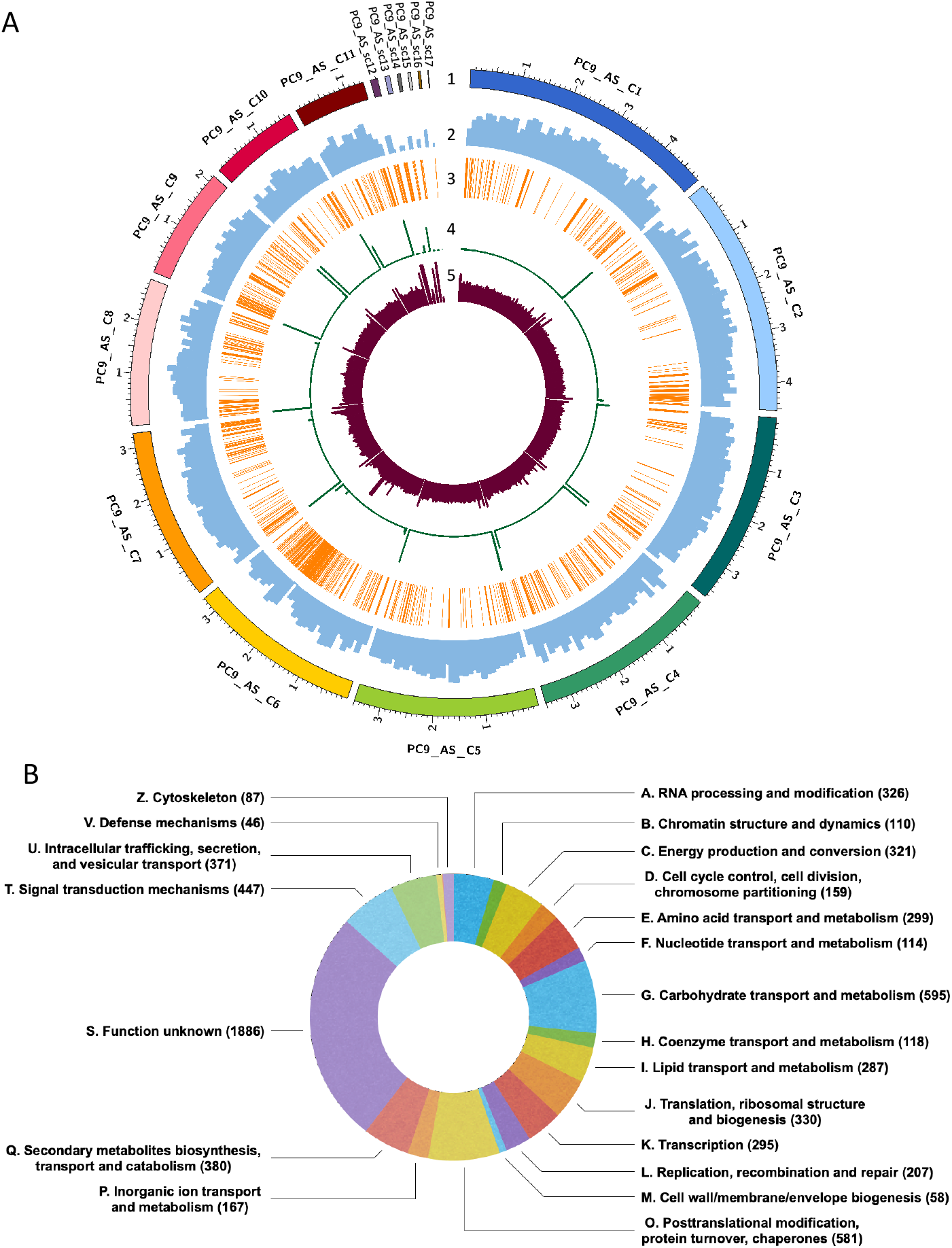
(A) Genome architecture of *P. ostreatus* strain PC9 based on our PC9_AS assembly. Tracks (outer to inner) represent the distribution of genomic features in our PC9_AS assembly: 1) sizes (in Mb) of PC9_AS scaffolds, with numbers prefixed by the letter “C” indicating the order of scaffold size; 2) gene density with 100-kb sliding windows, raging between 0-50 genes; 3) distribution of transposable elements along the PC9_AS genome; 4) distribution of telomere repeats with 10-kb sliding windows, ranging between 10-30 repeats; and 5) depth of gene coverage with 10-kb sliding windows, ranging between 0-300 depth. (B) Predicted functions of genes identified from the PC9_AS genome, cataloged using Cluster of Orthologous Groups (COGs) database.

The average coverage depth of PC9_AS is 135.3 reads per 10-kb (Figure 1A). Regions of high coverage depth are apparent at the ends of most scaffolds, perhaps due to the presence of telomeric repeats. Interestingly, regions of low coverage depth are found in some of the smallest scaffolds, such as 16 (45,074 bp) and 17 (9,086 bp). However, other small scaffolds show high coverage depth, such as 12 (135,750 bp), 14 (62,249 bp), and 15 (54,081 bp). The regions of higher coverage depth in these smaller scaffolds may correspond to the presence of repeats whereas, because of low repeat numbers in centromeric regions, the regions of lower coverage depth might represent misplaced centromeres. The gene density of PC9_AS in sliding windows of 100-kb is also illustrated in Figure 1A. Gene density is consistent among the different scaffolds, with an average of ∼33 genes per 100 kb, suggesting that aneuploidy does not occur in the PC9 genome. Moreover, regions of low gene density mostly align with stretches of telomeric repeats and with clusters of transposable elements (TEs). Telomeres consist of several tandemly arrayed (TTAGGG)n repeats, which hinder gene translation at these sites. Similarly, low expression of genes proximal to TEs has been reported previously (Castanera et al. 2016), potentially explaining low gene density in their vicinity. Although TEs are located near centromeres in certain fungi (Klein and O’Neill 2018), density of TEs in PC9_AS is highly diverse among most scaffolds (Figure 1A). A similar phenomenon has been reported for the previously published PC9_JGI and PC15 genomes (Castanera et al. 2016), wherein regions of low gene density aligned with zones harboring many TEs. We identified a total of 253 TE families in our PC9_AS genome, i.e., more than the 80 TE families reported previously for PC9_JGI and PC15 (Castanera et al. 2016). These TE families account for 7.12% of the total PC9_AS genome, greater than the 6.2% and 2.5% cited previously for the PC15 and PC9_JGI genomes, respectively (Castanera et al. 2016). This striking difference between PC9_AS and PC9_JGI may be due to the fragmented nature of the latter, a scenario that often hinders TEs identification. In Table 2, we summarize the transposable elements identified in PC9_AS, and it reveals that Class I transposons account for 89% of the all TEs, with LTR-retrotransposons being the most abundant family of Class I elements (Table S4). This outcome corroborates findings for the PC15 genome (Castanera et al. 2016). More detailed information on the TEs identified in PC9_AS genome—including size, number of fragments and repeats, and total bp—is presented in Table S4.

**Table 2.**
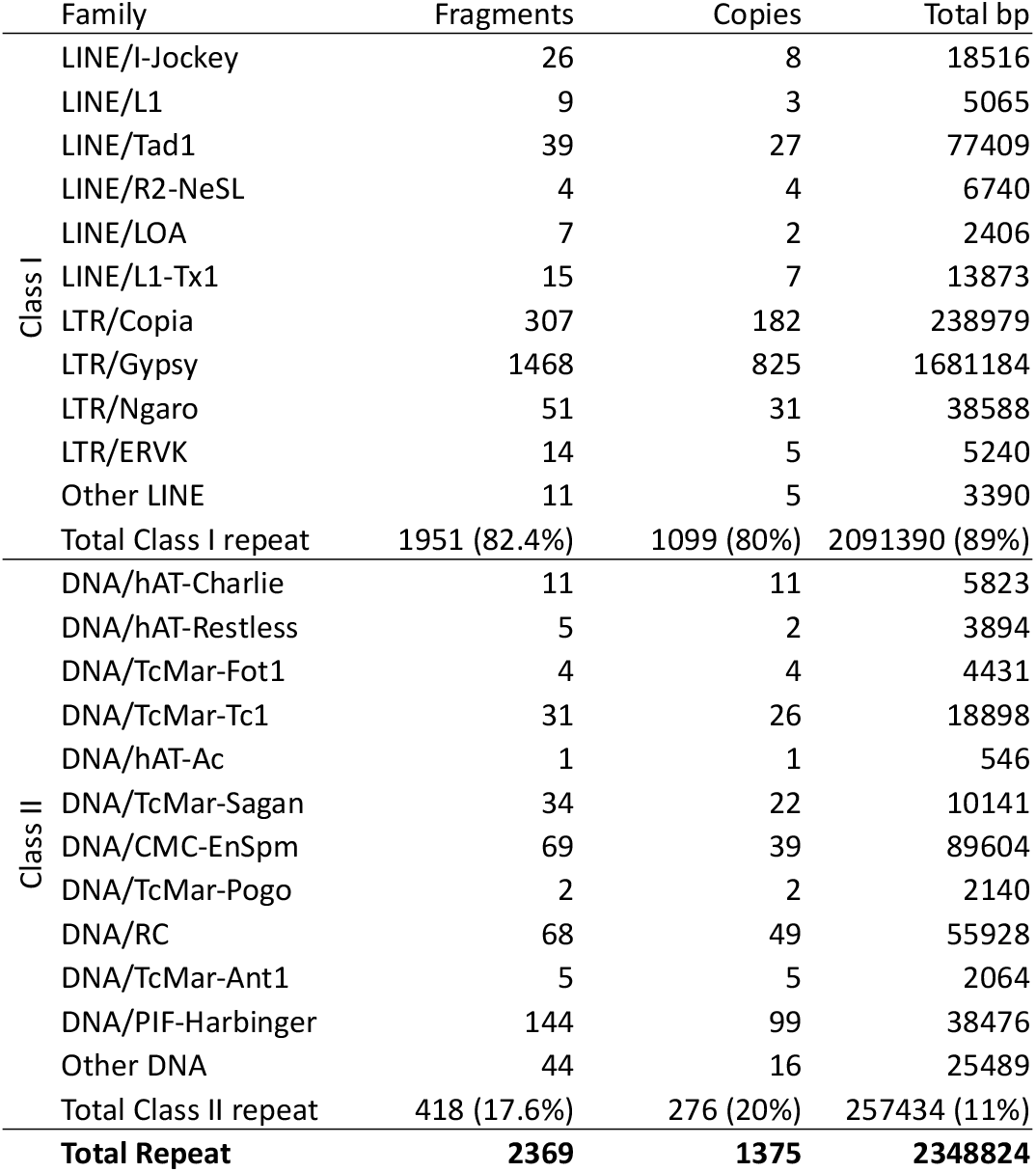
Classification of transposable elements identified from out *P. ostreatus* strain PC9_AS genome assembly.

|  | Family | Fragments | Copies | Total bp |
| --- | --- | --- | --- | --- |
| Class I | LINE/I-Jockey | 26 | 8 | 18516 |
|  | LINE/L1 | 9 | 3 | 5065 |
|  | LINE/Tad1 | 39 | 27 | 77409 |
|  | LINE/R2-NeSL | 4 | 4 | 6740 |
|  | LINE/LOA | 7 | 2 | 2406 |
|  | LINE/L1-Tx1 | 15 | 7 | 13873 |
|  | LTR/Copia | 307 | 182 | 238979 |
|  | LTR/Gypsy | 1468 | 825 | 1681184 |
|  | LTR/NGaro | 51 | 31 | 38588 |
|  | LTR/ERVk | 14 | 5 | 5240 |
|  | Other LINE | 11 | 5 | 3390 |
|  | Total Class I repeat | 1951 (82.4%) | 1099 (80%) | 2091390 (89%) |
| Class II | DNA/hAT-Charlie | 11 | 11 | 5823 |
|  | DNA/hAT-Restless | 5 | 2 | 3894 |
|  | DNA/TcMar-Fot1 | 4 | 4 | 4431 |
|  | DNA/TcMar-Tc1 | 31 | 26 | 18898 |
|  | DNA/hAT-Ac | 1 | 1 | 546 |
|  | DNA/TcMar-Sagan | 34 | 22 | 10141 |
|  | DNA/CMC-EnSpm | 69 | 39 | 89604 |
|  | DNA/TcMar-Pogo | 2 | 2 | 2140 |
|  | DNA/RC | 68 | 49 | 55928 |
|  | DNA/TcMar-Ant1 | 5 | 5 | 2064 |
|  | DNA/PIF-Harbinger | 144 | 99 | 38476 |
|  | Other DNA | 44 | 16 | 25489 |
|  | Total Class II repeat | 418 (17.6%) | 276 (20%) | 257434 (11%) |
|  | **Total Repeat** | **2369** | **1375** | **2348824** |

### Genome annotation

A total of 11,875 genes were annotated from PC9_AS, which is close to the current gene numbers of *P. ostreatus* PC15 and PC9_JGI genomes (12,330 genes and 12,206 genes, respectively). We used the Clusters of Orthologous Groups of proteins database (COGs) (Tatusov et al. 2000) to catalog the functions of those genes (Figure 1B). A considerable number of the gene functions are unknown, but the second biggest cluster in PC9_AS, containing 595 genes, consists of genes coding for putative carbohydrate transport and metabolism, such as transporters, chitinase, and carbohydrate-active enzymes (CAZymes), among others. CAZymes are essential to saprotrophic fungi like *P. ostreatus* to decay materials that are subsequently used as carbon sources (Mikiashvili et al. 2006). These enzymes play important roles in cellulose and hemicellulose degradation (Alfaro et al. 2016), with over 130 families described to date (Cantarel et al. 2009; Davies and Williams 2016), including glycosyl hydrolases (GHs), carbohydrate esterases (CEs), polysaccharide lyases (PLs), carbohydrate-binding modules (CBMs), glycosyl transferases (GTs) and auxiliary activities (AAs). We further explored the number of CAZymes-related genes in PC9_AS using the CAZymes database (Cantarel et al. 2009) and identified 459 such genes, an outcome consistent with our calculations for other *P. ostreatus* strains (Table S5; PC15=562; PC9_JGI=408). The EuKaryotic Orthologous Groups (KOG) database, which is a eukaryote-specific version of the COGs has been used previously to identify the orthologous and paralogous proteins from the PC15 and PC9_JGI genomes (Grigoriev et al. 2012). Therefore, we used the KOG classification available from the JGI for those genomes to reveal how gene families are distributed in *P. ostreatus* and to compare with our PC9_AS COG classification. As shown in Table S5, the biggest gene class among PC15 and PC9_JGI (both with 662 genes) is signal transduction mechanisms. However, PC9_AS only contains 447 genes coding for signal transduction mechanisms. In the classes of extracellular structures and nuclear structures, PC15 and PC9_JGI both have ∼100 genes associated with each class, whereas PC9_AS only harbors two extracellular structure genes and five nuclear structure genes. We acknowledge that these differences may reflect the use of different databases for gene function classification. Interestingly, PC9_AS also contained fewer PFAM domains (6770) compared to the other two *P. ostreatus* genomes (PC9_JGI=7914; PC15=7972).

### Genomic/genetic comparison of PC9 genomes

We performed a comprehensive comparison of the PC9_JGI (Alfaro et al. 2016; Castanera et al. 2016) and PC9_AS genomes. First, we observed that the number of scaffolds and contigs of PC9_AS (17) is smaller than that of PC9_JGI (572) (Table 1). In terms of size, both genomes are similar at ∼35 Mb, but PC9_JGI is slightly larger than PC9_AS by ∼600-kb. In terms of the N50 scaffold and N50 contig sizes, values for PC9_AS (N50 scaffold size = 3,500,734; N50 contig size = 3,500,734) are larger than those of PC9_JGI (N50 scaffold size = 2,086,289; N50 contig size = 99,058), which reflects greater contiguity in the former. PC9 and PC15 are protoclones of N001 (Larraya et al. 1999b), and the genome size of PC9 is slightly larger than that of PC15. The latest updated assembly of PC15 is distributed across 12 scaffolds, with a size of 34.3 Mb that is similar to PC9_AS. We used D-Genies (Cabanettes and Klopp 2018) to construct a dot-plot alignment of the genome between PC9_JGI and PC9_AS assemblies. The result, shown in Figure 2A, demonstrates that the PC9_JGI and PC9_AS genomes are highly similar, with small scaffolds of PC9_JGI corresponding to portions of the larger PC9_AS scaffolds, which indicates that our PC9_AS assembly is more complete. Moreover, when we aligned the two genomes in a circos plot (Figure 2B), we observed long regions of high similarity between the most relevant scaffolds of PC9_AS (scaffolds 1-11) (Table S6) and PC9_JGI (scaffolds 1-81) (Table S7). Figure 2B also reveals that, in general, more than five PC9_JGI scaffolds can be mapped to single PC9_AS scaffolds, with scaffolds 6 and 7 of PC9_AS incorporating 12 and 10 of the PC9_JGI scaffolds, respectively.

**Fig. 2.**
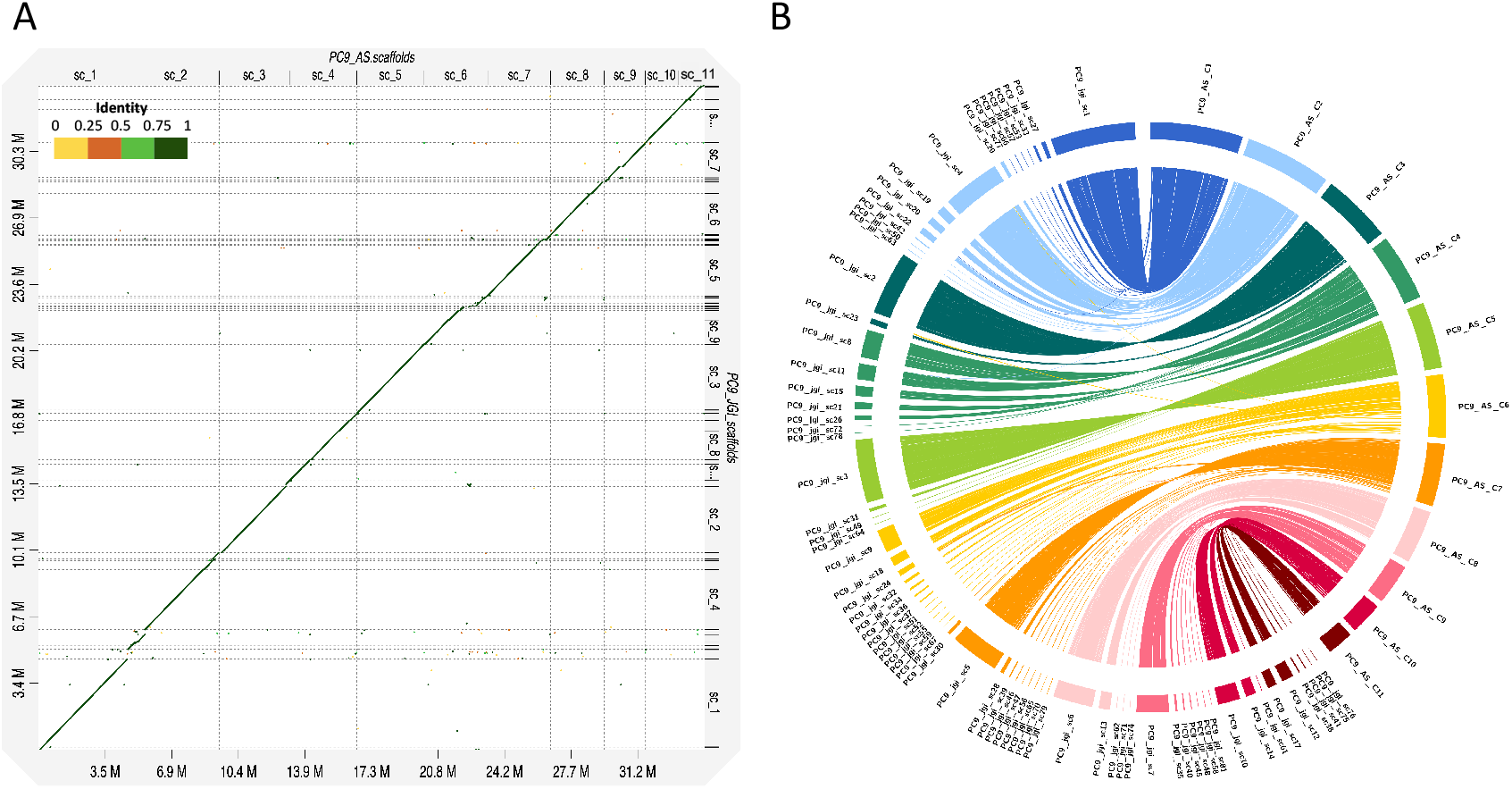
Comparisons between the PC9_AS and PC9_JGI *P. ostreatus* PC9 genomes. (A) Dot-plot alignment of *P. ostreatus* PC9_AS (target) and PC9_JGI (query) generated by D-Genies minimap2. (B) Circos plot showing regions of similarity shared between PC9_AS (scaffolds 1-11) and PC9_JGI (scaffolds 1-81) (identity>95%, length>10-kb).

## Conclusions

In this study, we used long PacBio and Illumina reads to assemble a new genome of *P. ostreatus* strain PC9. A combination of high read coverage and the latest bioinformatic tools resulted in a high-quality PC9_AS genome. Compared to the currently available PC9_JGI genome, our new assembly is more complete, comprising only 17 scaffolds that include six telomere-to-telomere scaffolds and the highest N50 values yet achieved for a *Pleurotus* genome. Genomic comparisons between PC9_AS and the current available assemblies of *P. ostreatus* evidence the high quality of our genome assembly. This new PC9 genome will enable the fungal research community to perform further genomic and genetic analyses of *P. ostreatus* and advance our understanding of this common edible mushroom.

## Acknowledgments

The authors thank Ursula Kües, Yoichi Honda, and Yitzhak Hadar for their helpful suggestions on growing *Pleurotus ostreatus*. We thank the IMB genomic core for Illumina sequencing. This work was supported by Academia Sinica Career Development Award AS-CDA-106-L03 and Taiwan Ministry of Science and Technology grant 106-2311-B-001-039-MY3 to YPH.

## Supplemental information

**Table S1.**
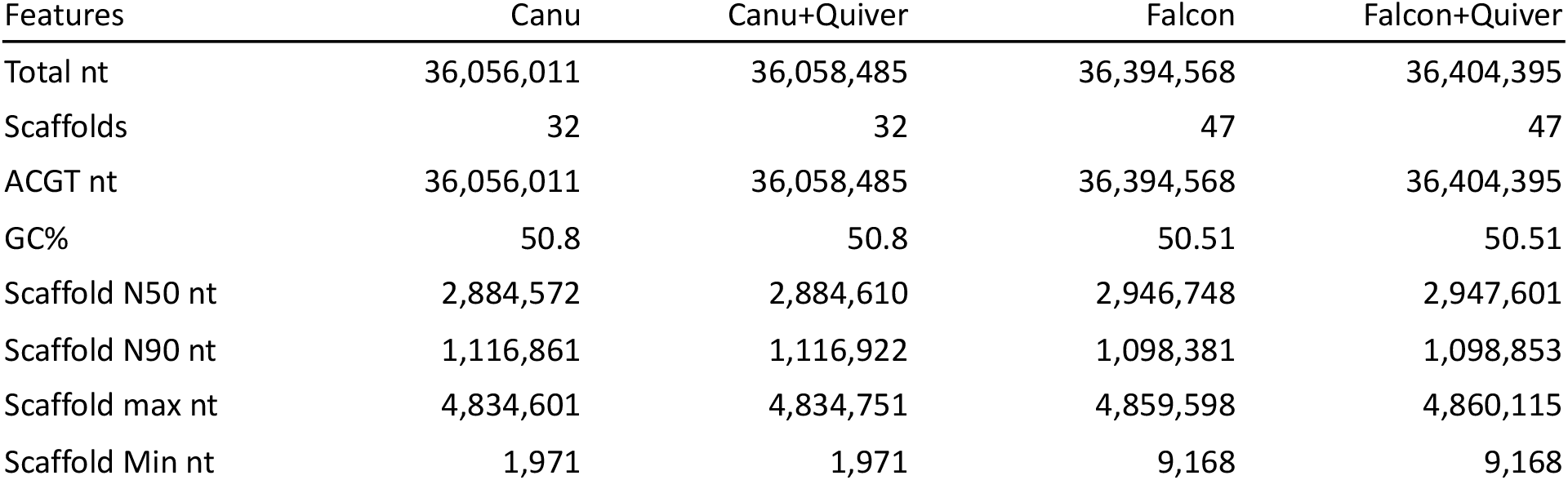
Features of the preliminary *P*.*ostreatus* PC9 assemblies obtained with Canu, Falcon, and Quiver (polishing). nt, nucleotides.

**Table S2.**
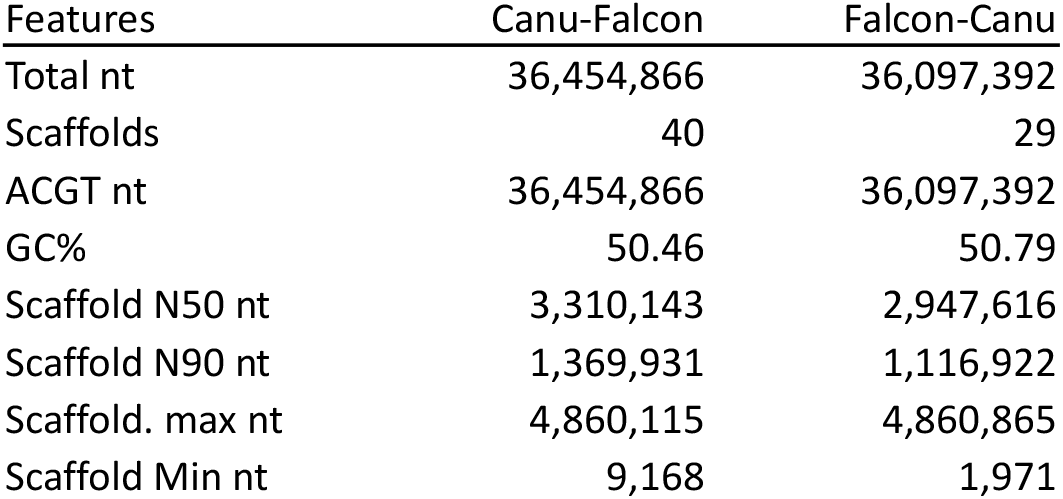
Features of the merged *P. ostreatus* PC9 assemblies constructed in Quickmerge, using the Canu assembly as reference and the Falcon assembly as donor (Canu-Falcon) or *vice versa* (Falcon-Canu). nt, nucleotides.

**Table S3.**
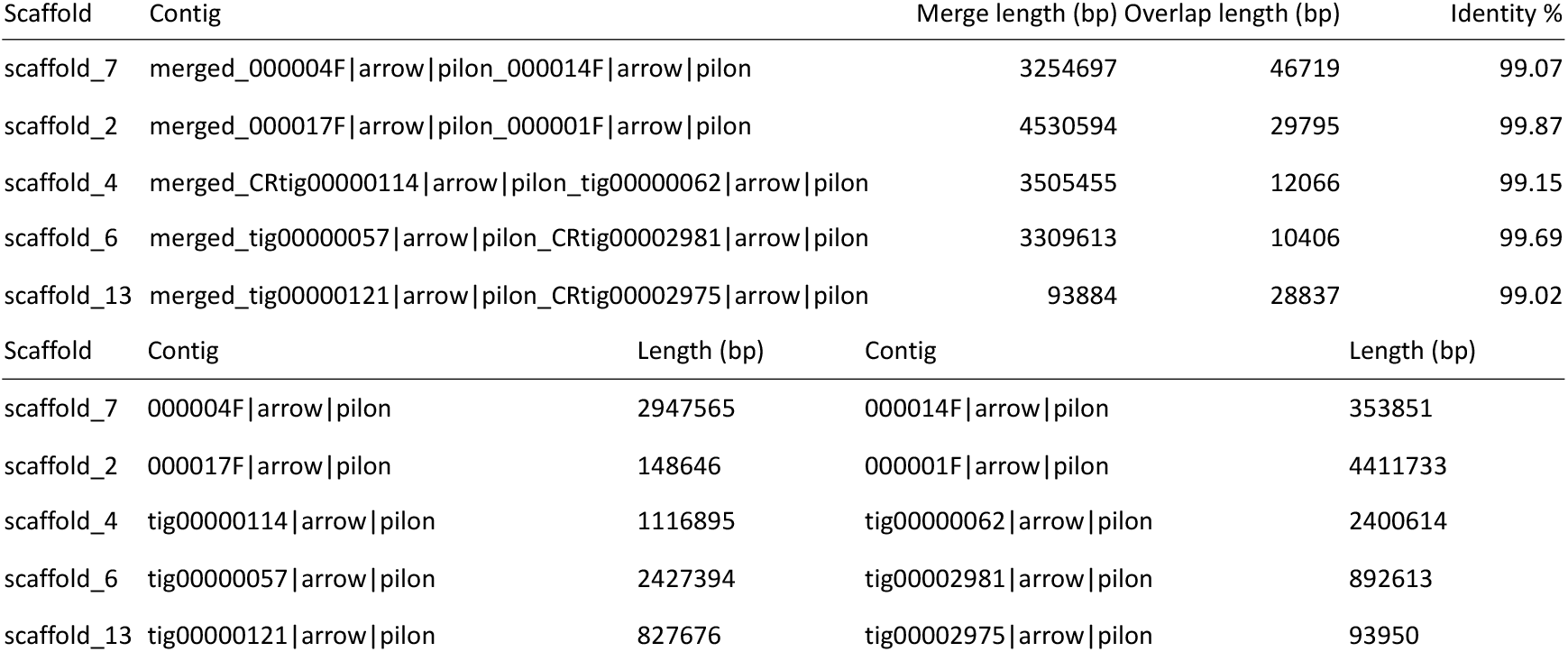
Information on 10 merged contigs of the PC9_AS assembly (identity>99%, overlap length>10-kb).

**Table S4.**
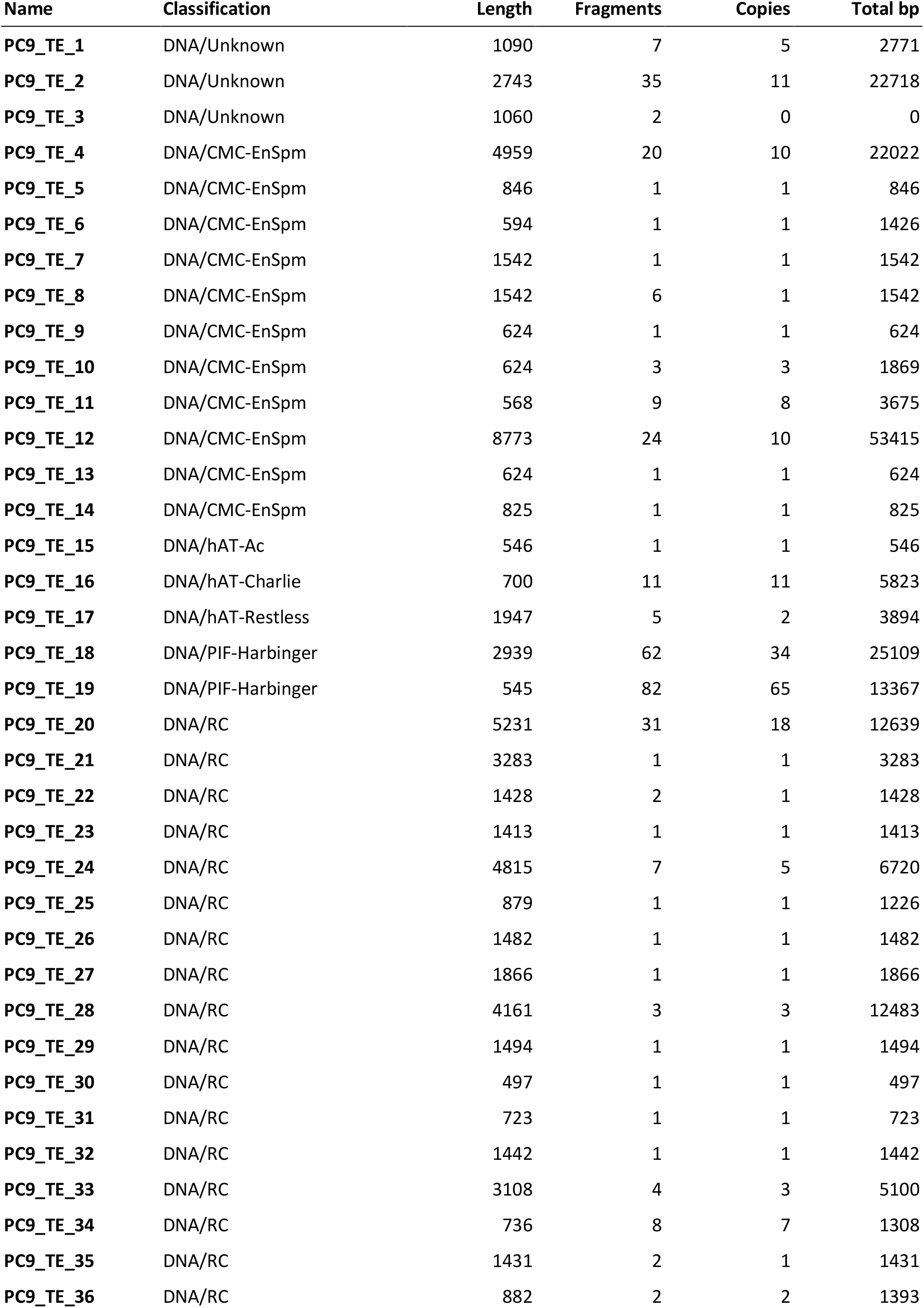
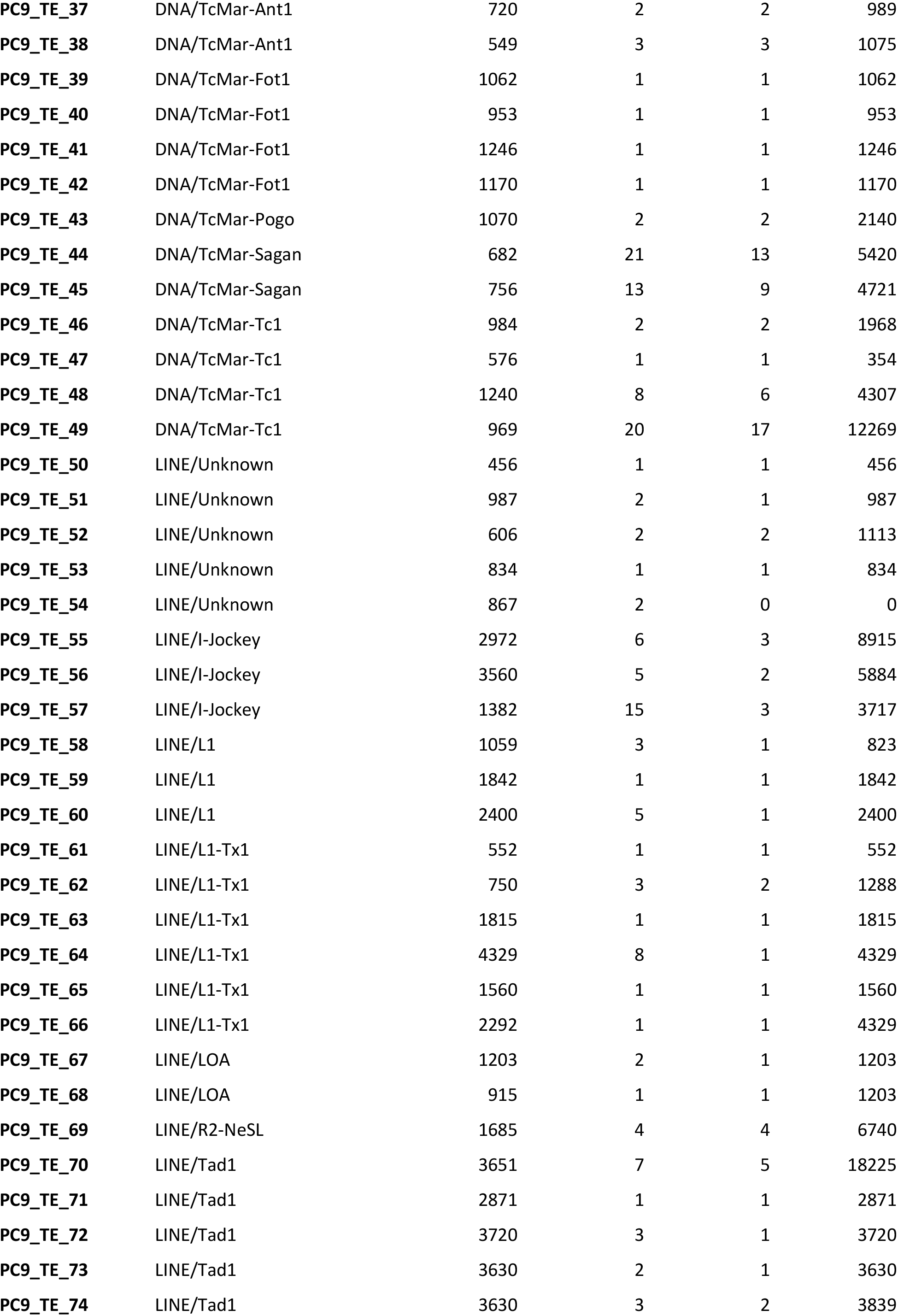
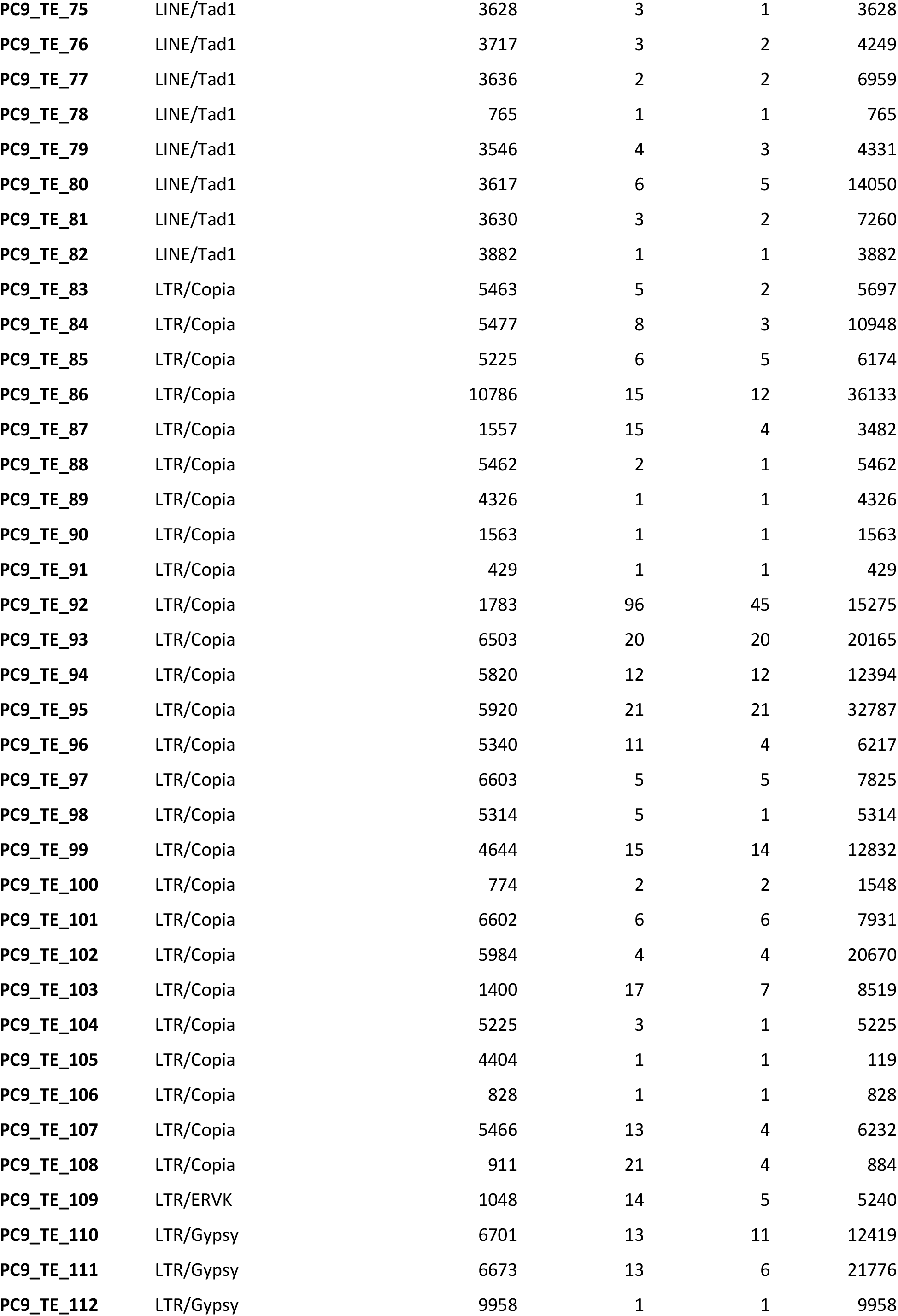
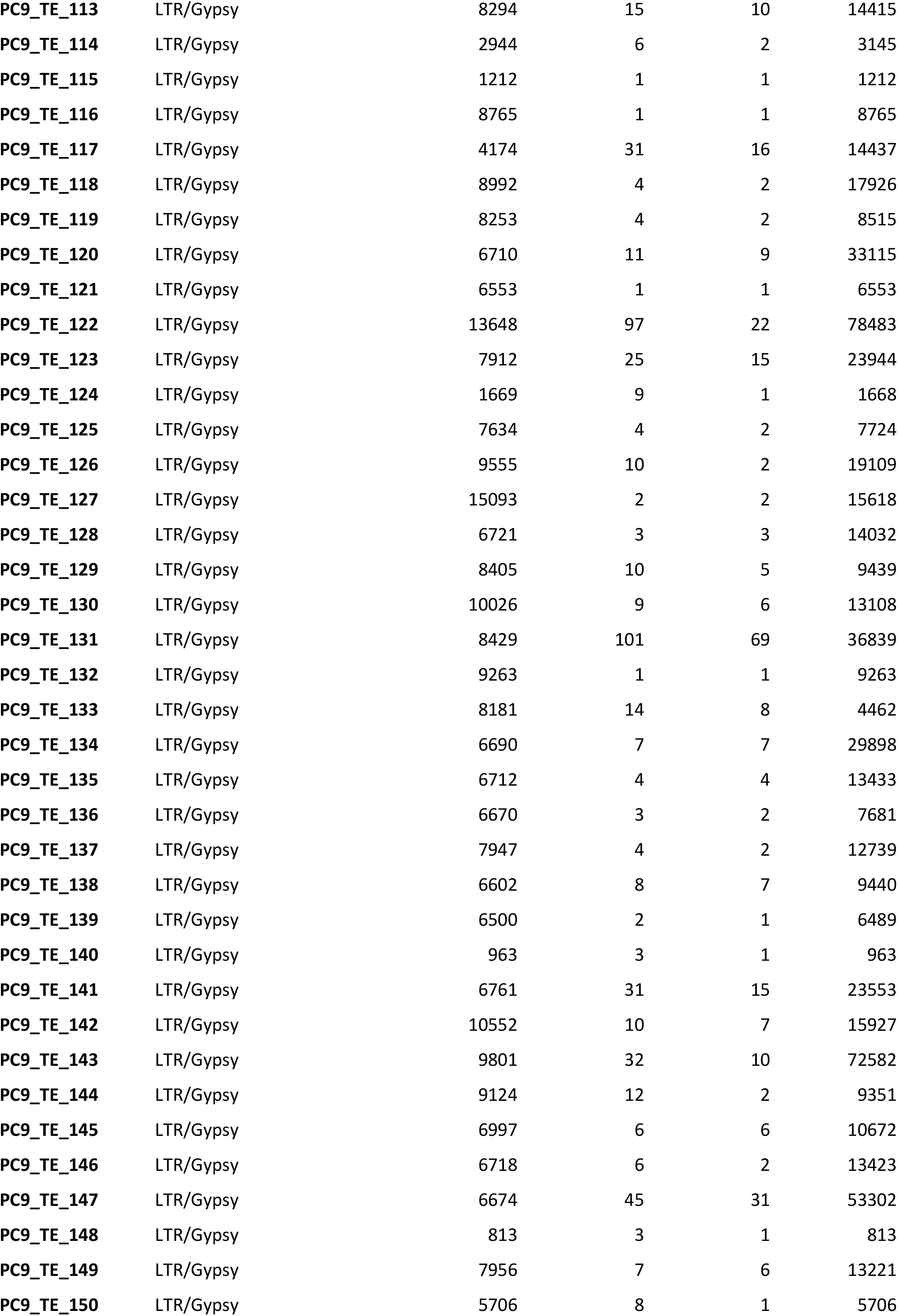
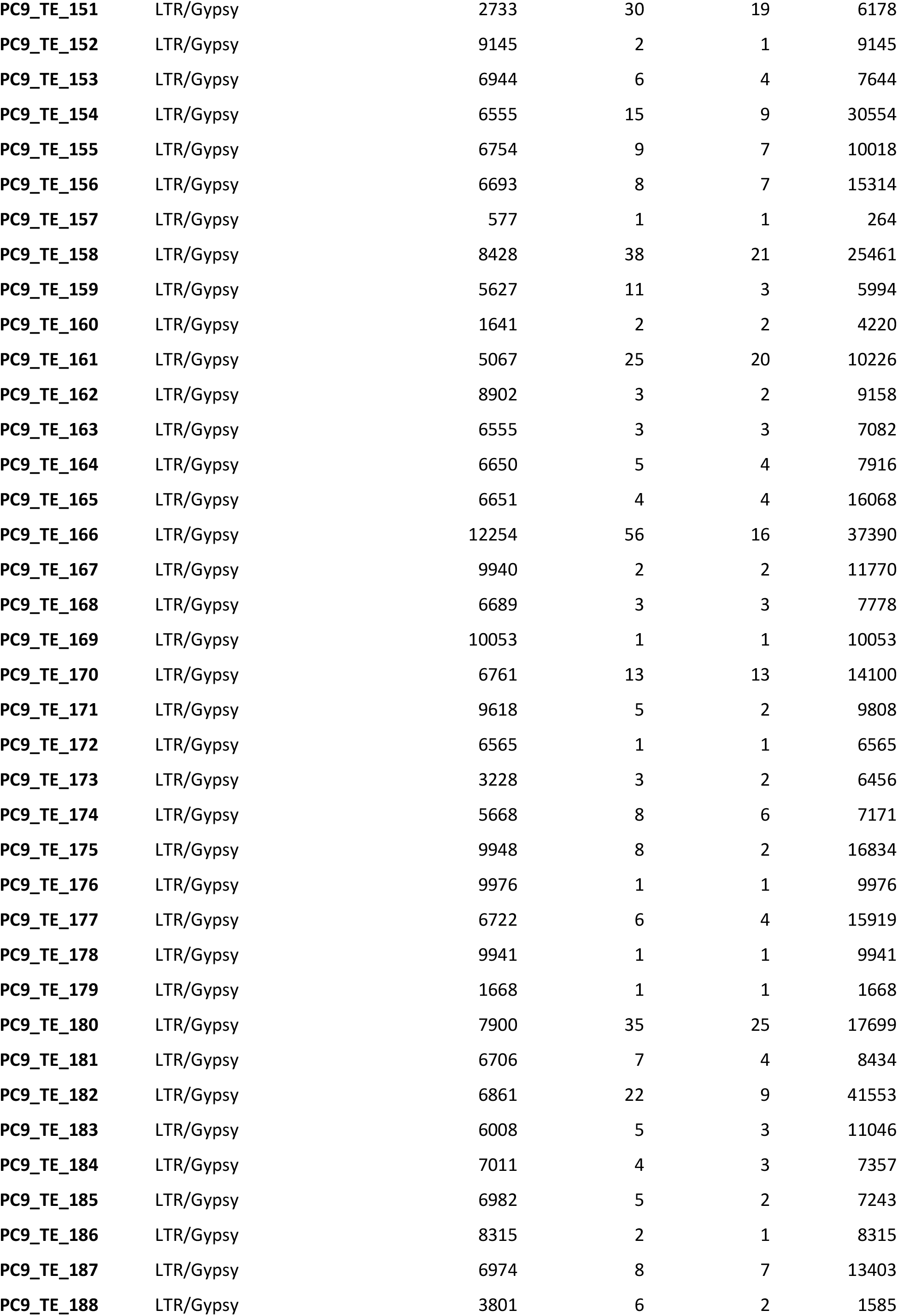
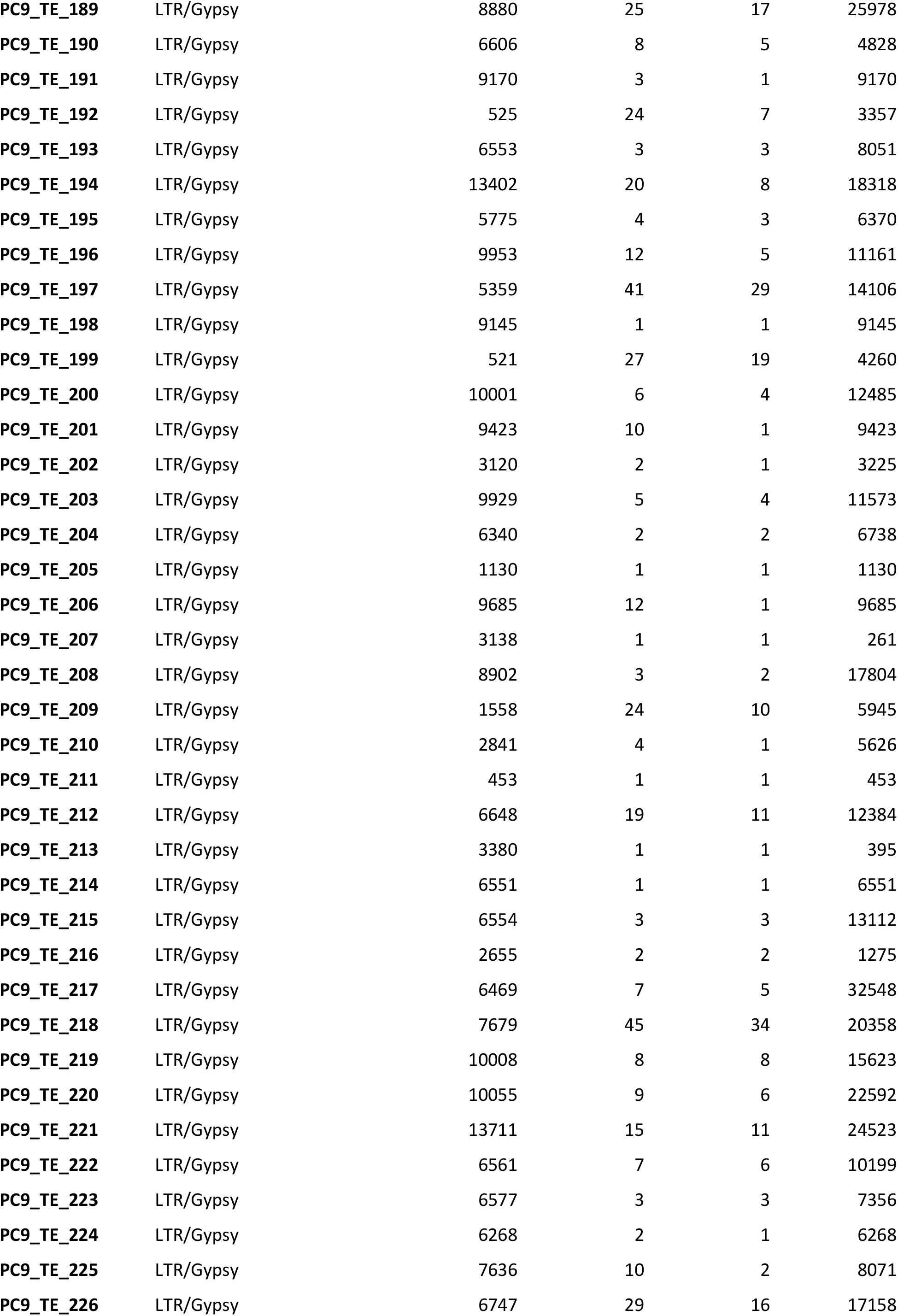
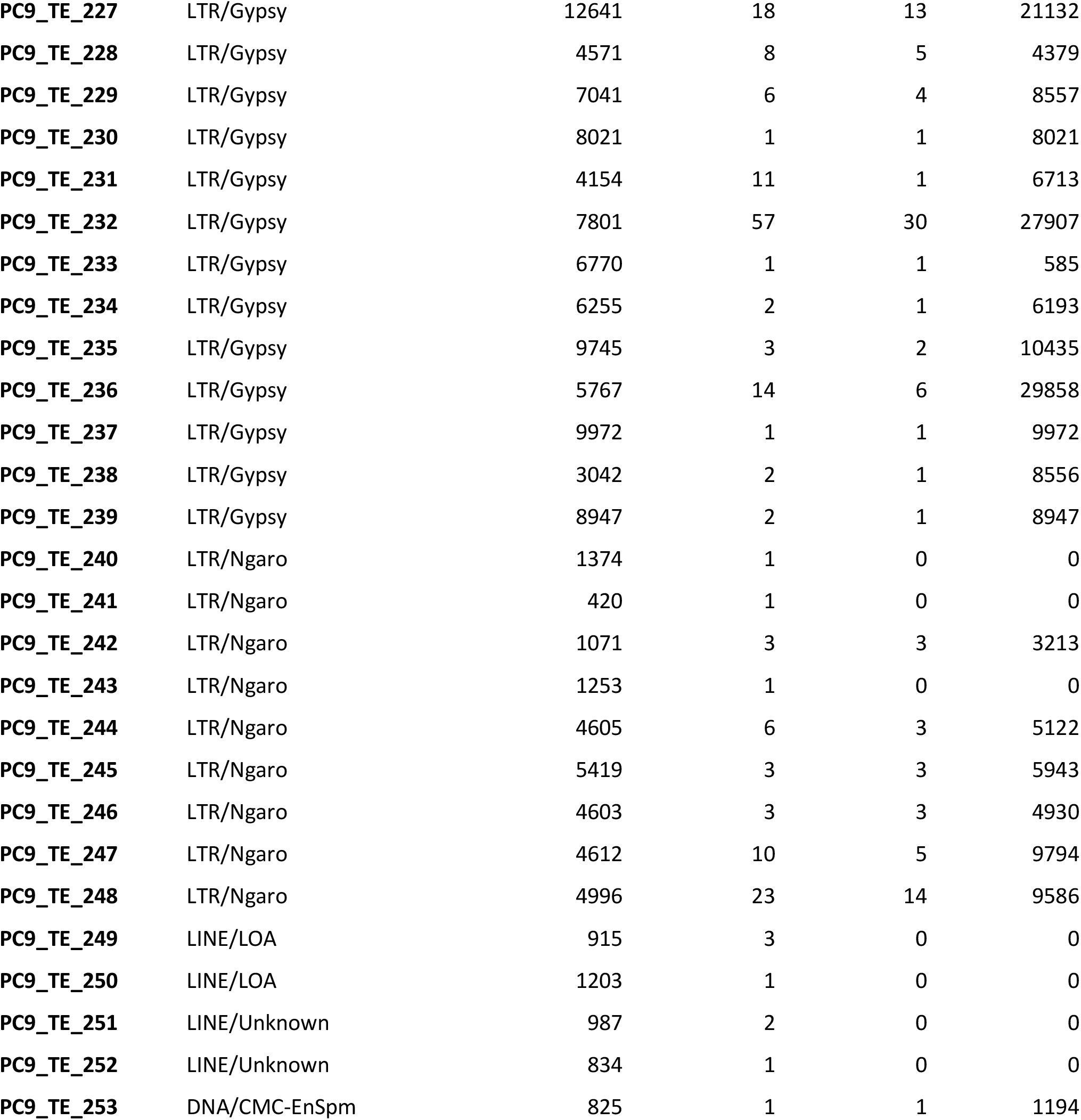
Families of transposable elements detected in the PC9_AS assembly.

**Table S5.** PFAM domains, CAZymes, and KOG/COG classifications for the PC15, PC9_JGI, and PC9_AS genome assemblies.

| Annotation | PC9_AS | PC9_JGI | PC15 |
| --- | --- | --- | --- |
| PFAMs | 6770 | 7914 | 7972 |
| CAZymes | 459 | 408 | 562 |
| KOG/COG classification | 7192 | 6321 | 6410 |
| A: RNA processing and modification | 326 | 373 | 381 |
| B: Chromatin structure and dynamics | 110 | 170 | 172 |
| C: Energy production and conversion | 321 | 305 | 312 |
| D: Cell cycle control, cell division, chromosome partitioning | 159 | 229 | 232 |
| E: Amino acid transport and metabolism | 299 | 243 | 243 |
| F: Nucleotide transport and metabolism | 114 | 80 | 82 |
| G: Carbohydrate transport and metabolism | 595 | 340 | 342 |
| H: Coenzyme transport and metabolism | 118 | 101 | 97 |
| I: Lipid transport and metabolism | 287 | 308 | 302 |
| J: Translation, ribosomal structure and biogenesis | 330 | 328 | 334 |
| K: Transcription | 295 | 427 | 439 |
| L: Replication, recombination and repair | 207 | 235 | 244 |
| M: Cell wall/membrane/envelope biogenesis | 58 | 72 | 78 |
| N: Cell motility | 1 | 6 | 6 |
| O: Posttranslational modification, protein turnover, chaperones | 581 | 659 | 651 |
| P: Inorganic ion transport and metabolism | 167 | 169 | 170 |
| Q: Secondary metabolites biosynthesis, transport and catabolism | 380 | 320 | 314 |
| S: Function unknown | 1886 | 407 | 432 |
| T: Signal transduction mechanisms | 447 | 662 | 662 |
| U: Intracellular trafficking, secretion, and vesicular transport | 371 | 335 | 337 |
| V: Defense mechanisms | 46 | 154 | 163 |
| W: Extracellular structures | 2 | 97 | 104 |
| Y: Nuclear structure | 5 | 110 | 100 |
| Z: Cytoskeleton | 87 | 191 | 213 |

**Table S6.** Scaffold sizes (in nucleotides) for our *P. ostreatus* PC9_AS assembly (scaffolds 1-17).

| Scaffold | Scaffold size (nt) |
| --- | --- |
| PC9_AS_sc1 | 4859873 |
| PC9_AS_sc2 | 4530689 |
| PC9_AS_sc3 | 3655511 |
| PC9_AS_sc4 | 3510511 |
| PC9_AS_sc5 | 3500734 |
| PC9_AS_sc6 | 3307864 |
| PC9_AS_sc7 | 3271080 |
| PC9_AS_sc8 | 2797324 |
| PC9_AS_sc9 | 2134864 |
| PC9_AS_sc10 | 1695225 |
| PC9_AS_sc11 | 1369937 |
| PC9_AS_sc12 | 135750 |
| PC9_AS_sc13 | 93126 |
| PC9_AS_sc14 | 62249 |
| PC9_AS_sc15 | 54081 |
| PC9_AS_sc16 | 45074 |
| PC9_AS_sc17 | 9086 |

**Table S7.** Scaffold sizes (in nucleotides) for the previously published *P. ostreatus* PC9_JGI genome (scaffolds 1-100; Alfaro et al. 2016).

| Scaffold | Scaffold size (nt) | Scaffold | Scaffold size (nt) | Scaffold | Scaffold size (nt) | Scaffold | Scaffold size (nt) |
| --- | --- | --- | --- | --- | --- | --- | --- |
| PC9_jgi_sc1 | 4430591 | PC9_jgi_sc26 | 232132 | PC9_jgi_sc51 | 15574 | PC9_jgi_sc76 | 10585 |
| PC9_jgi_sc2 | 3355223 | PC9_jgi_sc27 | 195390 | PC9_jgi_sc52 | 15315 | PC9_jgi_sc77 | 10582 |
| PC9_jgi_sc3 | 3327214 | PC9_jgi_sc28 | 179244 | PC9_jgi_sc53 | 15181 | PC9_jgi_sc78 | 10404 |
| PC9_jgi_sc4 | 3035439 | PC9_jgi_sc29 | 176908 | PC9_jgi_sc54 | 14718 | PC9_jgi_sc79 | 10219 |
| PC9_jgi_sc5 | 2563927 | PC9_jgi_sc30 | 162218 | PC9_jgi_sc55 | 14539 | PC9_jgi_sc80 | 10070 |
| PC9_jgi_sc6 | 2086289 | PC9_jgi_sc31 | 159002 | PC9_jgi_sc56 | 13955 | PC9_jgi_sc81 | 10029 |
| PC9_jgi_sc7 | 1693221 | PC9_jgi_sc32 | 135498 | PC9_jgi_sc57 | 13695 | PC9_jgi_sc82 | 9937 |
| PC9_jgi_sc8 | 1456251 | PC9_jgi_sc33 | 114947 | PC9_jgi_sc58 | 13616 | PC9_jgi_sc83 | 9854 |
| PC9_jgi_sc9 | 1215864 | PC9_jgi_sc34 | 99290 | PC9_jgi_sc59 | 13127 | PC9_jgi_sc84 | 9619 |
| PC9_jgi_sc10 | 1158825 | PC9_jgi_sc35 | 93408 | PC9_jgi_sc60 | 13062 | PC9_jgi_sc85 | 9584 |
| PC9_jgi_sc11 | 823721 | PC9_jgi_sc36 | 73952 | PC9_jgi_sc61 | 12936 | PC9_jgi_sc86 | 9566 |
| PC9_jgi_sc12 | 623273 | PC9_jgi_sc37 | 65148 | PC9_jgi_sc62 | 12473 | PC9_jgi_sc87 | 9545 |
| PC9_jgi_sc13 | 577815 | PC9_jgi_sc38 | 54882 | PC9_jgi_sc63 | 11699 | PC9_jgi_sc88 | 9468 |
| PC9_jgi_sc14 | 524972 | PC9_jgi_sc39 | 54229 | PC9_jgi_sc64 | 11564 | PC9_jgi_sc89 | 9372 |
| PC9_jgi_sc15 | 523842 | PC9_jgi_sc40 | 51136 | PC9_jgi_sc65 | 11483 | PC9_jgi_sc90 | 9295 |
| PC9_jgi_sc16 | 482463 | PC9_jgi_sc41 | 49577 | PC9_jgi_sc66 | 11391 | PC9_jgi_sc91 | 9279 |
| PC9_jgi_sc17 | 479568 | PC9_jgi_sc42 | 49327 | PC9_jgi_sc67 | 11281 | PC9_jgi_sc92 | 9173 |
| PC9_jgi_sc18 | 476552 | PC9_jgi_sc43 | 48231 | PC9_jgi_sc68 | 11143 | PC9_jgi_sc93 | 9065 |
| PC9_jgi_sc19 | 456527 | PC9_jgi_sc44 | 47637 | PC9_jgi_sc69 | 11128 | PC9_jgi_sc94 | 9029 |
| PC9_jgi_sc20 | 362764 | PC9_jgi_sc45 | 33979 | PC9_jgi_sc70 | 11031 | PC9_jgi_sc95 | 8970 |
| PC9_jgi_sc21 | 341089 | PC9_jgi_sc46 | 23569 | PC9_jgi_sc71 | 10991 | PC9_jgi_sc96 | 8956 |
| PC9_jgi_sc22 | 315898 | PC9_jgi_sc47 | 19536 | PC9_jgi_sc72 | 10945 | PC9_jgi_sc97 | 8840 |
| PC9_jgi_sc23 | 290907 | PC9_jgi_sc48 | 19420 | PC9_jgi_sc73 | 10890 | PC9_jgi_sc98 | 8812 |
| PC9_jgi_sc24 | 263270 | PC9_jgi_sc49 | 17084 | PC9_jgi_sc74 | 10838 | PC9_jgi_sc99 | 8526 |
| PC9_jgi_sc25 | 250080 | PC9_jgi_sc50 | 15720 | PC9_jgi_sc75 | 10613 | PC9_jgi_sc100 | 8476 |

